# Microbial functional pathways based on metatranscriptomic profiling enable effective saliva-based health assessments for precision wellness

**DOI:** 10.1101/2023.11.01.565122

**Authors:** Eric Patridge, Anmol Gorakshakar, Matthew M. Molusky, Oyetunji Ogundijo, Angel Janevski, Cristina Julian, Lan Hu, Momchilo Vuyisich, Guruduth Banavar

## Abstract

It is increasingly recognized that an important step towards improving overall health is to accurately measure biomarkers of health from the molecular activities prevalent in the oral cavity. We present a general methodology for computationally quantifying the activity of microbial functional pathways using metatranscriptomic data. We describe their implementation as a collection of eight oral pathway scores using a large salivary sample dataset (n=9,350), and we evaluate score associations with oropharyngeal disease phenotypes within an unseen independent cohort (n=14,129). As clinical validation, we show that the relevant oral pathway scores are significantly worse in individuals with periodontal disease, acid reflux, and nicotine addiction, compared with controls. Given these associations, we make the case to use these oral pathway scores to provide molecular health insights from simple, non-invasive saliva samples, and as molecular endpoints for actionable interventions to address the associated conditions.

**Highlights:**

- Microbial functional pathways in the oral cavity are quantified as eight oral scores
- Scores are significantly worse for individuals with oropharyngeal disease phenotypes
- This methodology may be generalized to other pathways and other sample types
- These scores provide longitudinal health insights in a precision wellness application

**Graphical Abstract:** 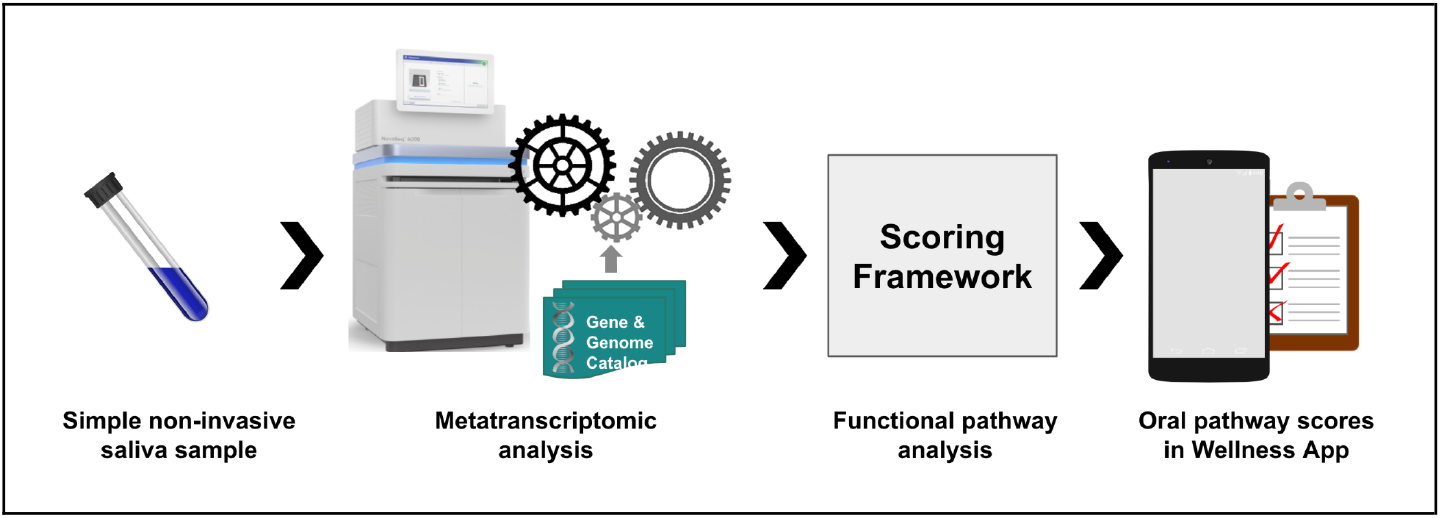

## 1. Introduction

Oral diseases, such as dental caries, periodontal gum disease, oral pre-malignancies such as leukoplakia, and oral cancers, impact billions of people [1,2]. The number of people affected by them continues to grow as a result of limited success of current oral health solutions such as fluoride [3,4], availability and affordability of food with high sugar content [5], lack of reliable oral cancer screening tools [6], and poor access to oral health care services in the community.

The oral cavity is the primary gateway to the body, and it hosts a complex environment which plays vital functional roles [8,9]. Both the oral microbiome and the oral immune system defend against a vast array of pathogens [10–13], but when either of these components is impaired, the oral cavity can support pathogenic activity, leading to chronic oral inflammation [14–17]. Since digestion begins in the oral cavity [18,19], the impairment of either component can lead to digestion-related health issues, including putrefaction of foods and host proteins within the mouth [20]. Halitosis (bad breath), gum disease, and oropharyngeal cancers are just a few of the conditions or diseases research suggests may coincide with an impaired oral microbiome and/or oral immune system [21–26].

Poor oral health may point to underlying health issues, since bi-directional associations exist between oral health and systemic health [27]. For example, left unchecked, chronic oral inflammation may advance to gum disease, and is linked to several conditions beyond the oral cavity, including diabetes, cardiovascular diseases, and Alzheimer’s disease [28–32]. Similarly, chronic halitosis is associated with Helicobacter pylori infections, liver disease, and gastroesophageal reflux disease [33–35]. Bi-directional associations like these are opportunities for health professionals to act on findings from early risk assessments and encourage proactive healthcare measures [36].

To improve oral, and indirectly, some aspects of systemic health, the existing prophylactic and therapeutic healthcare efforts can be augmented with non-invasive wellness tools that comprise molecular tests, diet changes, and supplement use [37,38]. As many have already shown, oral diagnostics are non-invasive tools to identify and address existing health issues [6], but these are reactive measures and generally not useful to proactive healthcare efforts [39–41]. In fact, there are still relatively few options which individuals or health professionals can leverage in the pursuit of proactive efforts for general healthcare [42].

We believe that an important step towards effective general healthcare is to provide deep health insights into the molecular activities prevalent in the oral cavity. Our recent efforts include a saliva-based test that measures the entire oral metatranscriptome [43] as well as the application of this test for personal wellness. A major aspect of our personalized precision wellness application is to provide each individual a detailed assessment of their molecular activities as a set of oral pathway scores, leveraging high-resolution detection of microbial functional features captured as KEGG Orthologs [44,45], or KOs, within the oral metatranscriptome. In the present context, we define an oral pathway score as a set of well-understood microbial biochemical functions that have been tied to various health conditions in the literature or clinical domain. The scope of the score and decisions to include or exclude specific functions are curated by domain experts using associations with various health conditions. The oral pathway scores we present in this paper were designed to assess common oral and systemic health issues, developed through a well-defined, data-driven process.

This paper aims to describe (1) the computational development methodology of saliva-based oral pathway scores using a large development cohort (n=9,350), and (2) their clinical validation against well-known oropharyngeal and related diseases within a large independent cohort (n=14,129). The oral pathway scores exclusively rely on the microbial KOs (i.e., microbial gene expression) measured in saliva samples, rather than on the taxonomy, since microbial functions are far more important for human health [46–49]. These functional scores enable us to understand the host-microbiome interactions in the oral cavity that maintain human health and can lead to development of methods that modulate microbial activities for improved health.

## 2. Materials and Methods

All samples and metadata were obtained from customers, at least 18 years old at the time of sample collection, who either completed a research Informed Consent Form (approved by a federally-accredited Institutional Review Board), or agreed to have their data analyzed in the terms and conditions, during the purchase of Viome’s Full Body Intelligence test. From each participant, we obtained a saliva sample, in addition to a comprehensive questionnaire describing their lifestyle, oral hygiene, dietary preferences, and health history. All study data are de-identified; data analysis team members have no access to personally identifiable information.

### 2.1. Cohort Development

The extensive collection of customer samples facilitates refreshing and rebuilding any reference cohort so that it reflects the customer population and molecular data, making each reference cohort a ‘living’ dynamic component for score development. The score development and validation cohorts described here are the latest iteration of such evolution.

To facilitate health insights from saliva samples, we design and validate wellness scores in the context of reference cohorts, each representative of the general adult customer population. Three guidelines are used to define these cohorts. First, we maximize the number of customer saliva samples available, and this serves as the initial snapshot for the cohorts. Second, around 10,000 of these samples are randomly selected (and this number is reduced later with specific criteria). Third, we examine the distribution of anthropometrics obtained through customer responses to questionnaires, and we exclude some outliers based on age, birth sex, and body mass index.

For the score development cohort, additional criteria serve to diminish artifacts related to sequencing depth or data sparseness; cohort samples contain a total count of more than 1,700 KOs, with a total read count of at least 70,000. The final reference cohort for score development is composed of 9,350 saliva samples from adults (59.4% female; see Table 1). For score validation, the final reference cohort is composed of 14,129 saliva samples from adults (62.9% female; see Table 1) that were not part of the score development framework (explained below).

**Table 1.** Representative anthropometric, sociodemographic, oral hygiene, and lifestyle metrics on reference cohorts. (Mean ± SD for continuous variables)

|  | Score Development Cohort<br>(n=9,350 saliva samples) | Independent Validation Cohort<br>(n=14,129 saliva samples) |
| --- | --- | --- |
| Female | 59.4% | 62.9% |
| Age (years) | 46.2 ± 13.0 | 46.0 ± 13.1 |
| BMI | 26.1 ± 5.4 | 25.8 ± 5.2 |
| Ethnicity | 74.2% White<br>From the remaining 25.8%:<br>28.1% Multi-ethnic<br>28.9% Hispanic or Latino<br>19.2% Asian<br>11.1% Black or African American<br>5.6% Arab or Middle Eastern<br>7.0% Other | 73.4% White<br>From the remaining 26.6%:<br>26.7% Multi-ethnic<br>29.1% Hispanic or Latino<br>19.9% Asian<br>10.6% Black or African American<br>6.7% Arab or Middle Eastern<br>6.9% Other |
| Tobacco use | 8.7% Current smoker<br>13.3% Former smoker<br>78.0% Never smoker | 8.3% Current smoker<br>13.4% Former smoker<br>78.2% Never smoker |
| Alcohol intake | 25.5% Heavy drinker<br>60.6% Light/Moderate drinker<br>9.2% Former drinker<br>4.5% Abstainer | 22.9% Heavy drinker<br>62.5% Light/Moderate drinker<br>9.8% Former drinker<br>4.7% Abstainer |
| Caffeine intake (drinks) | 1.5% High intake<br>19.2% Moderate intake<br>63.1% Low intake<br>16.1% Do not consume | 1.3% High intake<br>20.1% Moderate intake<br>62.2% Low intake<br>16.3% Do not consume |
| Mouthwash use | 23.8% Yes<br>75.9% No | 23.7% Yes<br>76.1% No |
| Brushing habit | 64.9% Two or more times a day<br>30.9% Once a day<br>4.0% Not every day | 67.1% Two or more times a day<br>29.3% Once a day<br>3.4% Not every day |
| Flossing habit | 9.0% Twice or more per day<br>35.5% Once a day<br>36.7% Less than once a day<br>18.6% Do not floss | 9.7% Twice or more per day<br>36.6% Once a day<br>35.6% Less than once a day<br>18.0% Do not floss |

### 2.2. Metatranscriptomic analysis

Saliva samples are collected and analyzed from individuals who fast and do not brush their teeth or use a mouthwash for at least 8 hours. The complete set of transcripts (RNA molecules) from each saliva sample is quantified using previously reported metatranscriptomic techniques [43], 3 yielding both the primary sequence and read count for each transcript. The bioinformatics method aligns each sequencing read to microbial genes, which are then grouped into KEGG Orthologs (KOs) using a custom catalog of genes and the KEGG databases [44]. Further details of the bioinformatics analyses can be found in Toma et al. [43]. The KOs we identify from saliva samples are used for downstream analyses including score development, validation, and eventually, scoring of new samples.

### 2.3. Score Development Framework

With the goal of delivering health insights based exclusively on microbial functions (i.e., gene expression) from the oral microbiome, we adopt an iterative and multistep process for developing wellness scores. All scores described in this paper comprise only microbial KOs. Within the finalized scores, each KO contributes a certain weight to the score, either in a positive or negative direction. To develop the scores, we examine domain concepts such as the activities associated with microbial colonization, consumption of salivary proteins, consumption of dietary mono and disaccharides, destruction of host tissues in the mouth, etc. We identify the microbial KOs associated with these physiological processes. The normalized expression value of each KO is aggregated with computationally determined weights applied to each, based on the first component of Principal Component Analysis (PCA). The final scores, when applied to a large cohort, exhibit a gaussian-like curve which stratify health insights across the population (shown in Supplemental Figure S1). As shown in Figure 1, for each score, the development steps include: 1) domain exploration to identify KOs and associated phenotypes; 2) metadata curation to enable case/control difference analyses; 3) signal definition to align each score for broad wellness assessment; 4) feature selection which involves iterative curation of score KOs; and 5) pathway activity quantification of the KOs selected for the score (Figure 1). This section explains each step in further detail using the score development cohort. It should be noted that even after finalizing a version of the scores, we continue to improve upon them while also advancing the multistep process.

**Figure 1.**
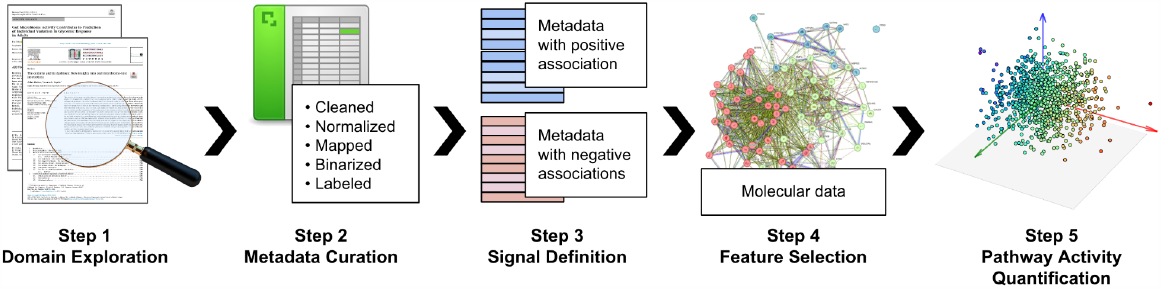
Score Development Framework.

#### Step 1: Domain Exploration

For each wellness score, we prioritize explainability of the overall score and its full set of KOs. We therefore begin by exploring the domain in order to understand the biological and clinical phenomena within the scope of each score, such as inflammation within periodontal pockets. While surveying knowledge sources, scientific manuscripts, and clinical literature, we collect an exhaustive set of KOs that are related to the score concepts and associated phenotypes, either based on their biological functions or based on reported correlations of their transcriptional regulation. Furthermore, iterating over this domain exploration step also allows us to prioritize a subset of KOs and phenotypes to be utilized in downstream development steps.

#### Step 2: Metadata Curation

Part of the score development process requires metadata labels for case/control difference analyses. To create these labels, we curate and normalize the phenotype data from customer surveys, including anthropometrics, socioeconomic, demographics, oral hygiene, and lifestyle characteristics. These sources are primarily self-reported and specifically include health conditions/diseases, which enable us to validate the association between these wellness scores and diseases. Other metadata is derived from sample data; for example, we label samples as “high cariogenic taxa” if they contained a comparatively high abundance of cariogenic species such as Streptococcus mutans. During the score development process, controls for each “case” definition are randomly sampled from the larger population, excluding anyone with a “case” label.

#### Step 3: Signal Definition

As we begin defining wellness scores for health insights, we use both the development cohort and the exhaustive set of curated KOs from Step 1. The metadata labels from Step 2 guide us during case/control difference analyses. Importantly, we make every effort to design scores for general wellness rather than diagnostics, so we avoid anchoring a score design to a single, individual disease. The definitive “signal” we pursue is a KO set that consistently differentiates related phenotypes (and other labels) across the development cohort. However, published gene expression efforts report variable findings across populations as well as reported measures [50–52], so it is not known which KOs will consistently differentiate signals across a large number of saliva samples. Therefore, we use a heuristic approach to prioritize KOs based on the numerous labels we create in Step 2. There are several metrics we utilize to define successful signals for phenotypes and health insights, and these include: 1) score difference between cases and controls; 2) a normalized score distribution; and 3) independence between score and sample sequencing depth. The output of the signal definition step is a minimal set of KOs, which sets a foundation for us to build upon for the final KO selection.

#### Step 4: Feature Selection

The final KO set for each score is defined through an iterative curation process, which uses all of the KOs and metadata labels identified through the above steps. We do not intentionally aim for each score to contain a specific number of KOs. Instead, iteration continues until scores reach an optimum, as determined through case/control difference analyses, stratifying individuals from the cohort into lower and higher scores, such that the metadata labels (phenotypes and other related labels) remain consistent. If the overall “signal” is not consistent, or the score distribution is far from a normal curve, or the score is highly correlated with sequencing depth, then the iterative process continues.

#### Step 5: Pathway activity quantification

Once the feature set (KOs) for a given pathway score is defined, the goal of this step is to combine the expression levels of the selected features to arrive at an aggregate quantification of the entire pathway. We derive a pathway score as a weighted function Score = C_1_F_1_ + C_2_F_2_ + … + C_n_F_n_, where F_i_ is the expression level of the feature and C_i_ is its weight. After experimenting with multiple weight computation methods, including a manual approach based on domain knowledge and decision tree methods, we settled on an algorithmic method that learns the weights based on the simultaneous expression of the selected features, which intuitively corresponds to the activity of the entire pathway. Since the first principal component of the principal component analysis (PCA) captures the largest variance in the original dataset, we believe that it provides a reasonable approximation of the simultaneous expression of the selected features. Furthermore, weighting features by the first principal component also enhances explainability of the score’s complete set of features.

### 2.4. Score validation framework

The validation cohort of 14,129 saliva samples is used to validate the reproducibility of the signals found in score development in an unseen independent sample set. To validate score designs, we ask whether scores can assess common oral health issues within the validation cohort. Domain knowledge provides evidence that many oral diseases increase risk for others: i.e., periodontal disease with halitosis [53]; periodontal disease with Sjögren’s [54]; acid reflux with both periodontal disease and cavities [55]. Therefore, we pursue a general wellness approach and examine the performance of all eight oral pathway scores with several diseases known to directly impact oropharyngeal health.

We specifically focus on three phenotypes – gingivitis, acid reflux, and nicotine addiction – for the validation process presented in this paper. To test scores with each disease phenotype, we perform case/control differential analyses. Case labels constructed for the development cohort are also applied to the validation cohort. Controls are defined as all individuals with no self-reported health conditions/diseases. Controls are matched to cases (1:10) on age, birth sex, and BMI. For age, we match individuals ≤ 10 year difference. For BMI, we match within BMI categories as follows: BMI < 18.5 (defined as ‘underweight’), 18.5 ≤ BMI < 24.9 (defined as ‘healthy’), 24.9 ≤ BMI < 29.9 (defined as ‘overweight’), 29.9 ≤ BMI (defined as ‘obese’). To perform case/control differential analyses, we use the Mann-Whitney U test along with the Benjamini-Hochberg correction to control the False Discovery Rate.

## 3. Results

### 3.1. Cohort characteristics

For score design and validation, we use a score development cohort and an independent validation cohort, respectively. The development cohort consists of 9,350 saliva samples from adults (59.4% Female) with an average age of 46.2 ± 13.0 and BMI of 26.1 ± 5.4. The validation cohort consists of 14,129 saliva samples from adults (62.9% Female) with an average age of 46.0 ± 13.1 and BMI of 25.8 ± 5.2. The large number of samples in each cohort is essential to identifying sufficient numbers of “case” metadata labels, which enable the case/control difference analyses presented herein.

Anthropometric, sociodemographic (including ethnicity), oral hygiene (including mouthwash use, brushing habits and flossing habits), and lifestyle characteristics (including tobacco use, alcohol intake, and caffeinated drink intake), are shown in Table 1 for the development and validation cohorts at the time of sample collection.

### 3.2. Metatranscriptomic analysis

Transcriptome metrics for score development (n=9,350) and validation cohorts (n=14,129) demonstrate the range and quality of sample data generated across both cohorts. Total single reads, or the number of reads in paired-end fastq file after demultiplexing, for the development and validation cohorts combined is 8.09 ± 3.34 in millions. The number of unique KOs (KO richness) associated with each sample for the development cohort and validation cohorts combined is 3,276 ± 488. The distribution for the number of unique species (Species richness) associated with each sample for the development cohort and validation cohorts combined is 349 ± 53.

### 3.3. Score Development

All eight oral pathway scores presented in this paper are shown in Table 2. Across these scores we include 234 distinct KOs, and we make every effort to minimize feature overlap between the scores; 221 of the KOs appear only once across all eight scores.

**Table 2.** High level overview of Oral Pathway Scores. Score names are shown alongside the high level descriptions of their scope, and the number of KOs in each score. In a pairwise comparison of each score, there is a maximum of two KOs shared between any two scores.

| Score Name | Key functions | Microbial activities | KOs |
| --- | --- | --- | --- |
| *OralAmmoniaProductionPathways* | Ammonia production<br>Arginine biosynthesis<br>Nitrogen metabolism | Homeostatic levels of ammonia within the oral cavity which protect against acids and impart an anti-cariogenic effect. | 26 |
| *OralButyrateProductionPathways* | Butanoate products<br>Lysine degradation<br>Short chain fatty acid metabolism | Production of beneficial butyrate as well as butyrate from pathogens. | 30 |
| *OralCariogenicPathways* | Acid production<br>Chelation activity<br>Sucrose metabolism | Cariogenic and chelation activities, as well as other activities which promote dental cavities. | 28 |
| *OralFlagellarAssemblyPathways* | Bacterial secretion system<br>Chemotaxis<br>Flagellar assembly<br>Quorum sensing | Flagellar components, related motility, informative of pathogenic activity. | 49 |
| *OralLPSBiosynthesisPathways* | Biofilm formation<br>Biosynthesis of nucleotide sugars<br>Lipopolysaccharide biosynthesis | Production of pro-inflammatory lipopolysaccharide (LPS). | 35 |
| *OralPeriodontalMucin-DegradationPathways* | Amino sugar metabolism<br>Galactose metabolism<br>Glycoprotein metabolism<br>Nucleotide sugar metabolism | Degradation of salivary mucin and diminishment of oral health barrier defenses. | 30 |
| *OralPeriodontalPro-InflammatoryPathways* | Glyoxylate metabolism<br>Lipoic acid metabolism<br>Porphyrin metabolism<br>Protease/peptidase activity | Inflammatory factors and degradation of periodontal gum health. | 26 |
| *OralUreaseActivityPathways* | Acid-stress<br>Nickel transport<br>Urate metabolism<br>Urea metabolism | Acid-stress and pH balancing activities as well as others which protect against an acidic environment and related pathogenic activity. | 29 |

As detailed in section 2.3 of the Materials and Methods, score development is a five-step process that begins with domain exploration to identify microbial KOs potentially relevant to the score concepts and associated phenotypes. Here, we detail the score development process using OralUreaseActivityPathways as an illustrative example. (Each of the other scores follow a similar process.)

As part of domain exploration, we identify key components and functions that we consider to be “in scope” for the *OralUreaseActivityPathways* score (Table 3), which focuses on the oral microbiome’s acid/base activities related to urea.

**Table 3.** Scope of the *OralUreaseActivityPathways* Score.

|  |  |  |
| --- | --- | --- |
| **Key components** | Organic acids | Citrate, succinate, fumarate, 2-oxoglutarate |
|  | Proteins | Urease, urea carboxylase, urea transporter, organic acid transporter, stress regulator, argininosuccinate lyase |
|  | Purines | Uric acid, xanthine, hypoxanthine, adenine, guanine |
|  | Enzyme cofactors | Nickel, nickel-pincer cofactor, manganese, iron |
|  | Amino acids | Arginine, citrulline, ornithine, glutamine, glutamate |
|  | Other | Urea, allantoate, ureidoglycine, N-acetylglutamate, Acetyl-CoA, ammonia, nitrate, nitrite |
| **Key functions** | Nitrogen waste | Urea metabolism, urate metabolism, ammonia production |
|  | Signaling | Acid-stress, acid tolerance |
|  | Homeostasis | Protection, carbon metabolism |
|  | Other | pH balance, nutrient deprivation |

After extensive research covering the scope of the score, we identify a long list of relevant KOs (in the hundreds or thousands), and we supplement these with additional relevant KOs which we identify from the development cohort. At this point in the process, all KOs are thought to directly or indirectly contribute to a “key component” or “key function” shown in Table 3.

The second and third steps of the Score Development Framework involve metadata curation and signal definition (see section 2.3 of the Materials and Methods). The metadata labels we consider to be central to the OralUreaseActivityPathways score include Dry mouth, Kidney disease, Kidney cyst, and Sjögren’s syndrome, from a total of more than 500 labels available.

The fourth step is feature selection, and the output of this step is the final set of KOs and their weights (loadings) to the score. Using OralUreaseActivityPathways as an example, Table 4 shows the included KOs and their respective weights. As presented here, the KOs with positive loadings are indicative of beneficial contributions with respect to the oral microbiome’s acid/base status and related to urea.

**Table 4.** Score design for *OralUreaseActivityPathways*. Loadings originate from the first principal component of PCA analysis using the development cohort.

| Loadings | KEGG Ortholog (KO) and Description |
| --- | --- |
| 0.23282 | K00101 (lldD); L-lactate dehydrogenase (cytochrome) |
| 0.21168 | K03746 (hns); DNA-binding protein H-NS |
| 0.20788 | K01420 (fnr); CRP/FNR family transcriptional regulator |
| 0.20710 | K11605 (sitC); manganese/iron transport system permease |
| 0.17742 | K11606 (sitD); manganese/iron transport system permease |
| 0.10082 | K00605 (gcvT,AMT); aminomethyltransferase |
| -0.00871 | K01895 (ACSS1_2,acs); acetyl-CoA synthetase |
| -0.01382 | K15585 (nikB,cntB); nickel transport system permease |
| -0.01632 | K15586 (nikC,cntC); nickel transport system permease |
| -0.03862 | K01941 (uca); urea carboxylase |
| -0.04997 | K03187 (ureE); urease accessory protein |
| -0.07264 | K07770 (cssR); two-component OmpR, response regulator |
| -0.09710 | K01755 (argH,ASL); argininosuccinate lyase |
| -0.10985 | K01218 (gmuG); mannan endo-1,4-beta-mannosidase |
| -0.13493 | K07650 (cssS); two-component OmpR, histidine kinase |
| -0.14145 | K14977 (ylbA,UGHY); (S)-ureidoglycine aminohydrolase |
| -0.15198 | K01466 (allB); allantoinase |
| -0.17054 | K02083 (allC); allantoate deiminase |
| -0.17658 | K01428 (ureC); urease subunit alpha |
| -0.18129 | K01430 (ureA); urease subunit gamma |
| -0.20003 | K09477 (citT); citrate:succinate antiporter |
| -0.21234 | K01429 (ureB); urease subunit beta |
| -0.21740 | K03190 (ureD,ureH); urease accessory protein |
| -0.22403 | K03188 (ureF); urease accessory protein |
| -0.26017 | K14048 (ureAB); urease subunit gamma/beta |
| -0.26711 | K00366 (nirA); ferredoxin-nitrite reductase |
| -0.27764 | K22373 (larA); lactate racemase |
| -0.29448 | K00370 (narG,narZ,nxrA); nitrate reductase, alpha subunit |
| -0.31953 | K03191 (ureI); acid-activated urea channel |

### 3.4. Score validation

We set out to determine whether there is a significant association between scores and disease phenotypes within an unseen independent cohort. Using the validation cohort, matched case/control difference analyses are conducted with oropharyngeal diseases, demonstrating that the final scores are able to differentiate phenotypes across a set of samples independent from those we use in score development.

Three oropharyngeal and related disease phenotypes are used to validate oral pathway scores, matching for case/control difference analyses (Figure 2). Cases of gingivitis (n=150; c=1,462), cases of acid reflux (n=450; c=4,268), and cases of nicotine addiction (n=67; c=665) are all matched (1:10). Control samples are selected from those lacking all disease phenotype labels in the validation cohort. As seen in Figure 2, the differences between “case” and “control” samples are variably significant across the eight scores (p ≤ 0.05 to p ≤ 0.001) according to the Mann-Whitney U test with Benjamini-Hochberg correction for multiple hypothesis testing (FDR < 0.05).

**Figure 2.**
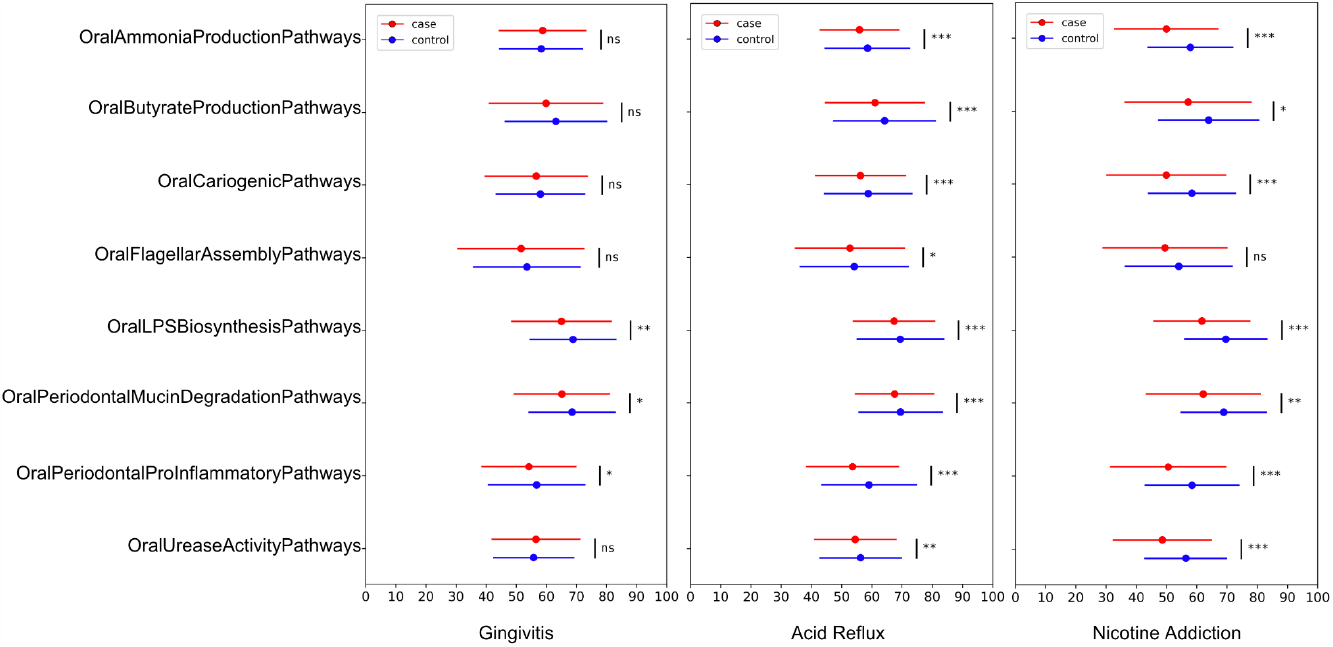
Oropharyngeal and related diseases negatively impact oral scores. Oral scores from our independent validation cohort comparing people with self-reported oropharyngeal and related diseases (case in red) versus people with no self-reported comorbidities (control in blue). Cases include gingivitis, acid reflux, and nicotine addiction (as indicated). Cases and controls are matched (1:10) by age, sex, and BMI. The color line represents mean ± SD of each score in case and control respectively; *p ≤ 0.05, **p ≤ 0.01, ***p ≤ 0.001, ns p > 0.05, from the Mann-Whitney U test with Benjamini-Hochberg correction for multiple hypothesis testing (FDR < 0.05).

## 4. Discussion

Oral microbiomes are associated with both oral and systemic diseases. Therefore, our starting point in this study was that molecular data obtained from the oral microbiome may provide useful biomarkers of health and disease. Further, it is clear from multiple studies [56,57] that the functional aspects of the microbiome are likely more impactful on human physiology than composition (taxonomy), which only measures the functional potential. Our metatranscriptomic method quantifies gene expression, that is the activity of microbial functions, and makes them available as KEGG Orthologs or KOs, which we use as the basis of developing oral functional pathway scores.

In the design of our oral pathway scores, both the development and independent validation cohorts are considerably important to the process. A comparison of the two cohorts (Table 1) suggests they are similar and comparable in terms of anthropometrics, sociodemographics, oral hygiene (including mouthwash use, brushing habits and flossing habits), and lifestyle metrics (including tobacco use, alcohol intake, and caffeinated drink intake), supporting our goal of validating the reproducibility of the signals found in score development within an unseen independent validation cohort.

While developing each oral pathway score, we prioritize health insights for oropharyngeal disease phenotypes, leveraging the connection between the oral microbiome and oral health. The key functions we identify for each score (Table 2) are relevant to disease phenotypes, and these functions take central roles in the score designs and guide the selection of specific KOs. In the case of *OralUreaseActivityPathways*, the expanded set of key functions (Table 3) point to several urea-related KOs which are up-regulated at low pH or during acid stress, and these guide the design for this score. While finalizing each KO set, we minimize correlations with the sequencing depth (r=0.08 for *OralUreaseActivityPathways*) so that scores are robust (less sensitive to the amount of data generated). We also work towards a normal distribution of scores across the development cohort (See Supplemental Figure S1) to maximize their utility as wellness scores. The case/control difference analyses with oropharyngeal and related disease phenotypes (Figure 2) indicate these wellness scores are effective for oral health insights, and this signal is reproducible in a very large cohort of independent samples.

The choice of the eight oral pathway scores presented here was based on the biochemical and biological functions that underlie the disease phenotypes we focused on for this study – gingivitis, acid reflux, and nicotine addiction – which were available to us in a large validation cohort. While we focus on a few health conditions, the development and validation methodology presented here is general enough to quantify a large array of functional pathways. Indeed, we are currently in the process of developing and validating not only the full oral microbiome, but also the gut microbiome and the human blood transcriptome scores.

Our approach to validating oral pathway scores is to test the significance of the score in a case cohort with the disease phenotype of interest, compared with a healthy control cohort without that disease phenotype. The key question in this study, of course, is whether a signal found during score development is reproducible in independent collections of samples. Both the case and control validation cohorts come from a sample set that was independent and unseen at score development time. We recognize that there could be other ways to validate the mechanistic basis of these pathways, for example via independent laboratory analysis of the resulting metabolite measurements for each pathway – this could be possible future work.

Among cases of gingivitis, three of the expected oral pathway scores demonstrated significant differences with respect to controls (Figure 2), *OralLPSBiosynthesisPathways, OralPeriodontalMucinDegradationPathways*, and *OralPeriodontalProInflammatoryPathways*. These scores were designed to assess microbial activities which strongly align with symptoms of gingivitis (Table 2). In its earliest form, before progressing to full periodontal disease, gingivitis symptoms include irritation, redness, and swelling of the gums (inflammation), although these are reversible with proper care and maintenance. The *OralPeriodontalMucinDegradationPathways* score is focused on salivary barrier defenses, which preserve the health of the oral cavity, and mucin degradation is a known factor in periodontal disease [58–61]. The design of the *OralPeriodontalProInflammatoryPathways* score is heavily focused on both inflammation and periodontal destruction, which are defining symptoms of gingivitis and periodontal disease [62–64]. The *OralLPSBiosynthesisPathways* score overlaps with proinflammatory signals, but it is designed to typify inflammatory activities, specifically highlighting contributions of a detrimental Gram-negative biofilm [65–68]. As gingivitis develops into full periodontal disease, we anticipate other scores would gain significance, and we look forward to examining this in future work.

In looking at cases of acid reflux, all eight of the oral pathway scores demonstrated significant differences with respect to controls (Figure 2). These results are consistent with an oral environment associated with acid reflux [69,70]. In addition, mounting evidence indicates that acid reflux (and the low pH of gastric acid) diminishes both dental health [71–73] and oral soft tissue health [74–76]. Reports also show disease associations increase with gastric acid contact [77,78]. Three of the oral pathway scores, *OralAmmonia-ProductionPathways, OralCariogenicPathways*, and *OralUreaseActivityPathways*, are strongly aligned with the acidic environment fostered by the low pH of gastric acid. And, as already indicated, several scores are strongly aligned with gingival damage and inflammation. In future studies, we look forward to examining how interventions designed to address the low pH and/or contact time of gastric acid will impact the scores.

We also found significant differences for seven of the oral pathway scores between cases of nicotine addiction and controls (Figure 2), and these results are consistent with an oral environment disrupted by tobacco products [79,80]. Among cases (individuals who self-reported a nicotine addiction) and controls, 92.5% and 8.4% were current tobacco users, respectively. Tobacco smoke contains over four thousand compounds, most of which are considered toxic in both gaseous and solid form [81,82], and nicotine itself is known to impact many systems across the human body [83–85]. With regards to oral health, it is well-established that tobacco products have a detrimental impact on every oral condition as well as the success of many oral treatments [86]. Specifically, it has been shown that tobacco products can diminish salivary pH [87,88], reduce salivary flow [88,89], and exacerbate bone loss and inflammation [90–92]. Long term tobacco use is also known to increase risk for dental caries [93], periodontitis [94], oral cancer [95], and many other oral conditions [96,97]. Thus, the overall impact of tobacco-related oral conditions aligns with the design and performance of our scores.

Besides affecting oral health, there is increasing evidence that the oral microbiome is linked to systemic health. Our extensive catalog of over 500 systemic disease phenotype labels affords us the opportunity to explore connections to systemic health as an additional aspect of oral pathway scores. For example, with regards to the *OralUreaseActivityPathways* score, there are well-established connections between salivary urea levels and kidney function [99,100]. Therefore, during score development, we prioritize phenotypes like Kidney disease and Kidney cyst, which retain high explainability with the *OralUreaseActivityPathways* score. Given this, we can do a case/control differential analysis for *OralUreaseActivityPathways* in our independent validation cohort. Here, “case” samples are defined using an aggregate of multiple systemic disease phenotypes which are based on reported associations to urea or uric acid levels in saliva (or blood) [55,56], including: Alcohol addiction [57,58], Ankylosing spondylitis [59], Diverticulosis [60], Gout [56], Hyperlipidemia [61,62], Kidney cyst [63,64], Kidney disease [65–67], Pancreatitis [68,69]. “Control” samples are selected from those lacking all disease phenotype labels in the validation cohort. We did such an initial analysis, with case (n=388) and control (n=3,633) matched (1:10) by age, sex, and BMI. The result was a significant difference in score values (p ≤ 0.001) using the Mann-Whitney U test with Benjamini-Hochberg correction for multiple hypothesis testing (FDR < 0.05). Similar analyses could be done for each score presented here, but this is outside the scope of this paper, and will be attempted as future work.

The study presented here has some limitations. Our validation employs case/control difference analyses which do not account for all possible confounders. Our analyses do not evaluate causality; the KO features that make up the scores are most likely a combination of causal and consequential. Prospective interventional trials will identify the causal features in follow-on studies.. While our cohorts are very large and include many demographics, they may not perfectly represent every group in the USA or other countries. Finally, our metadata labels are based on self-reported data that may be misreported; however, given the size of our cohorts, such effects should be negligible.

One important objective of this work is to develop an iterative process for continuously improving the design of these scores (the selection of features and their weights). We will continue to improve the utility of the scores as we collect more data from more samples and individuals, and as the field of molecular science advances to include additional insights. The goal for each score is to provide the most meaningful health insights given the latest science and data.

Another important future objective of this work is to show the impact of lifestyle factors and specific interventions on functional pathways, over the course of a longitudinal re-testing regime. For example, there are several interventions for periodontal gum disease that could be initiated by an individual as part of a self-care routine, or by a dental professional who sees acute or chronic issues. An important question is how these oral pathway scores change longitudinally from a pre-intervention state to a post-intervention state. We look forward to evaluating the effect of a range of interventions on molecular functional pathway scores in the future.

## 5. Conclusions

With the increasing prevalence of many types of chronic diseases due to lifestyle factors, and the advent of broadly available molecular testing, it is important to establish a systematic methodology for assessing microbial functional pathways relevant to these diseases. Chronic diseases being complex and multifactorial, it is necessary for such a methodology to assess a group of related molecular markers together (rather than individual molecular markers). In this paper, we have presented such a methodology, in the context of the oral microbiome providing functional pathway insights associated with oropharyngeal disease phenotypes. We have described the development of eight oral functional pathway scores using a large cohort, and shown that the scores are able to distinguish oropharyngeal disease phenotype cases from controls in an unseen independent validation cohort. The same methodology can be applied to other contexts, such as the gut microbiome providing functional pathway insights associated with gastrointestinal and related disease phenotypes.

To our knowledge, these are the first reported wellness scores based on the oral metatranscriptome, and they are designed to provide molecular health insights from simple, non-invasive saliva samples. In this context, it is possible to aggregate many oral scores as presented here, into an overall score that represents oral health at the highest level; indeed, this is the approach we have taken in our implementation within our wellness product. The product also shows the longitudinal changes to the scores at multiple time points, for individuals who re-test their oral health. Furthermore, the health insights delivered by these oral pathway scores can drive hygiene, dietary, lifestyle, and pharmaceutical recommendations [98].

## CRediT authorship contribution statement

**Eric Patridge:** Conceptualization, Data curation, Investigation, Formal Analysis, Methodology, Software, Visualization, Writing – original draft, and Writing – review & editing. **Anmol Gorakshakar:** Data curation, Investigation, Formal Analysis, Methodology, Software, Visualization, Writing – original draft, and Writing – review & editing. **Matthew M. Molusky:** Data curation, Investigation, Formal Analysis, Methodology, Software, Visualization, Writing – original draft, and Writing – review & editing. **Oyetunji Ogundijo:** Data curation, Formal analysis, Software. **Angel Janevski:** Software. **Cristina Julian:** Investigation, Validation, Writing - review & editing. **Lan Hu:** Investigation, Software, Writing - review & editing. **Momchilo Vuyisich:** Funding acquisition, Investigation, Resources, and Writing – review & editing. **Guruduth Banavar:** Conceptualization, Funding acquisition, Investigation, Project administration, Resources, Supervision, Writing – original draft, and Writing – review & editing.

## Data availability

The raw data used in this study cannot be shared publicly due to privacy and legal reasons. However, if data is specifically requested, we may be able to share a summary and/or portions of the data. Researchers requiring more data for non-commercial purposes can request via: https://www.viomelifesciences.com/data-access. Viome may provide access to summary statistics through a Data Transfer Agreement that protects the privacy of participants’ data.

## Declaration of Generative AI and AI-assisted technologies in the writing process

No AI-assisted technology was used in the writing process.

## Declaration of Competing Interest

The material represents original research, has not been previously published (is deposited as a preprint in **biorxiv.org**) and has not been submitted for publication elsewhere while under consideration in CSBJ. All authors of this manuscript were employees of Viome Life Sciences Inc at the time of their contributions, and held stock options in the company. MV and GB hold management positions within the company. The work was funded by Viome Life Sciences Inc.

## Supplementary data

**Figure S1.**
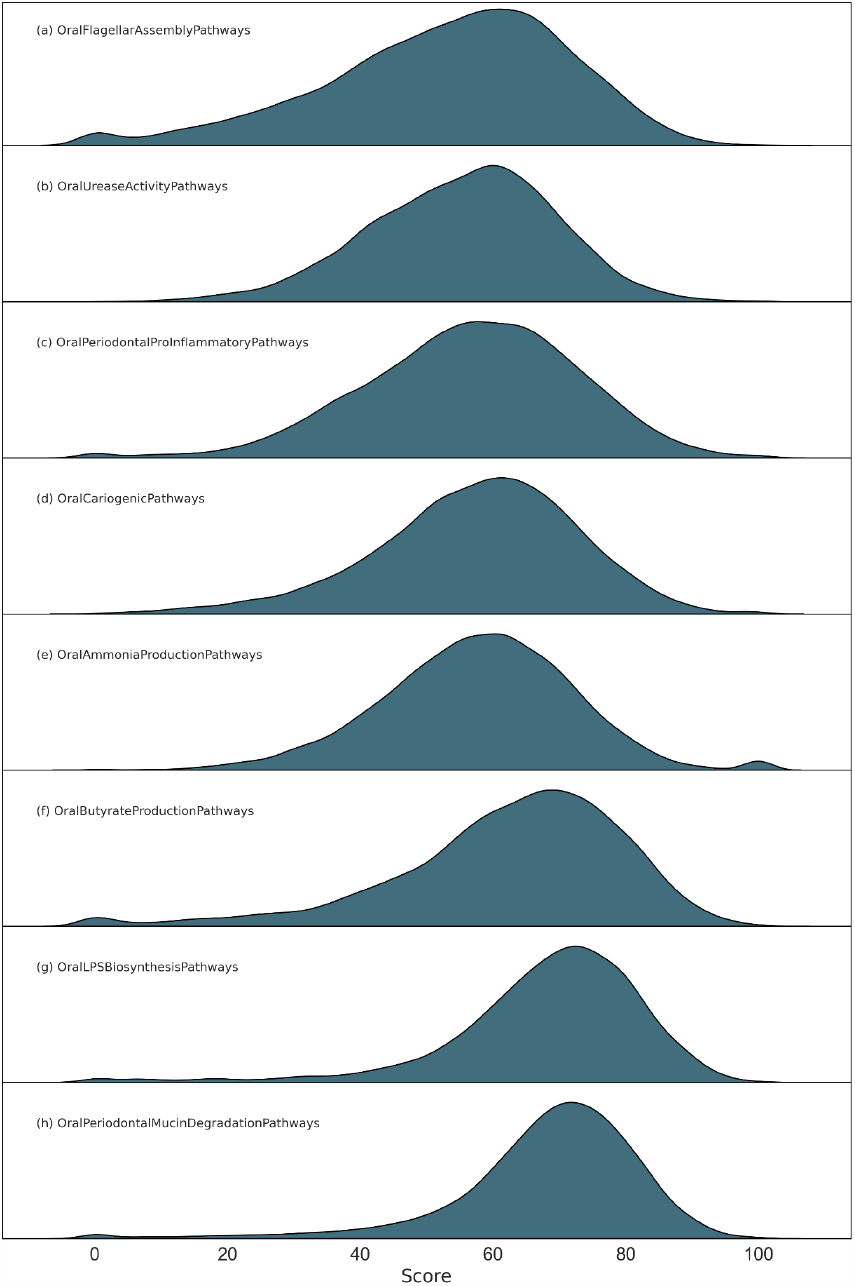
Landscape of score distribution. The distribution of eight functional pathway scores in an independent cohort (n=14,129 samples). The functional score, represented as Principal Component 1 (PC1) rescaled to the development cohort for each sample, is expected to follow a Gaussian normal distribution. Some scores deviate significantly from this normal distribution, which can be attributed to missing molecular signatures in the reference cohort, resulting in extremely high or low scores. Scores exceeding 100 or falling below 0 were clipped to 100 or 0, respectively. The secondary peaks observed in scores a, c, e, f, g, h are artifacts resulting from score clipping.

